# Bayesian nonparametric (non-)renewal processes for analyzing neural spike train variability

**DOI:** 10.1101/2023.10.15.562381

**Authors:** David Liu, Máté Lengyel

## Abstract

Neural spiking activity is generally variable, non-stationary, and exhibits complex dependencies on covariates, such as sensory input or behavior. These dependencies have been proposed to be signatures of specific computations, and so characterizing them with quantitative rigor is critical for understanding neural computations. Approaches based on point processes provide a principled statistical framework for modeling neural spiking activity. However, currently, they only allow the instantaneous mean, but not the instantaneous variability, of responses to depend on covariates. To resolve this limitation, we propose a scalable Bayesian approach generalizing modulated renewal processes using sparse variational Gaussian processes. We leverage pathwise conditioning for computing nonparametric priors over conditional interspike interval distributions and rely on automatic relevance determination to detect lagging interspike interval dependencies beyond renewal order. After systematically validating our method on synthetic data, we apply it to two foundational datasets of animal navigation: head direction cells in freely moving mice and hippocampal place cells in rats running along a linear track. Our model exhibits competitive or better predictive power compared to state-of-the-art baselines, and outperforms them in terms of capturing interspike interval statistics. These results confirm the importance of modeling *covariate-dependent* spiking variability, and further analyses of our fitted models reveal rich patterns of variability modulation beyond the temporal resolution of flexible count-based approaches.

## 1 Introduction

Analyses of spike time [49, 76, 86] and count [12, 26, 57] statistics have revealed neural responses *in vivo* to be structured but generally variable or noisy [24, 44, 70, 78]. To model this stochastic aspect of neural spike trains, probabilistic approaches based on temporal point processes have been widely applied. This in turn has been a major driver of point process theory development [42] for capturing spiking variability structure with statistical models [74]. The study of neural computation underlying naturalistic behavior in particular involves non-stationary spike trains, which presents a significant challenge as apparent spiking variability is a result of both irreducible “intrinsic” neural stochasticity as well as dependencies on behavioral covariates that can themselves vary on multiple time scales.

Different approaches have been proposed for handling non-stationary spike trains, starting with the classical log Cox Gaussian process [14, 55] to allow variations in the local intensity or firing rate while modeling independent spikes. Dependencies on previous spikes can be captured to first order with renewal processes, and these models have been extended to non-stationary cases through modulation of the hazard function with some time-dependent function [43, 79] or through rescaling interspike intervals with a covariate-dependent rate function [2, 6]. Another approach based on Hawkes processes and spike-history filters [45, 83, 91] introduces conditional point processes that go beyond the first order Markov assumption. Approaches based on recurrent networks [51, 90] and neural ODEs [10, 41] can in theory capture arbitrarily long dependencies on past spikes and input covariates, but provide more limited descriptive interpretability.

However, not only the rate but also the variability of spiking encodes task-relevant information [34, 60], and bears signatures of the underlying computations [11]. Importantly, this variability has stimulus- and state-dependent structure [12, 21, 48, 67]. In statistical modeling language, this corresponds to heteroscedastic or input-dependent observation noise. Such structure reflects computations performed in the underlying neural circuit, and thus characterizing it from data in a flexible and robust manner is critical for advancing theories of neural computation. The classical approaches reviewed above do not attempt to characterize such covariate-dependent changes in variability. Flexible count models have been introduced to more faithfully capture variability at the count level [28], and recent work has extended this to the general case of input-dependent variability [48]. Count approaches however are limited in the resolution of the analysis set by the time bin size. In addition, the resulting count statistics strongly depend on the chosen time bin size [48, 73].

While firing rates are routinely modeled as input-dependent, extending point process models with input-dependent variability has not been widely explored in the literature. Rate-rescaled and modulated renewal processes rely on fixed base renewal densities. Allowing the shape parameters of the renewal density to vary with covariates in the corresponding hazard function is one potential approach, but this still relies on a commitment to a particular parametric family of renewal densities. Spike-history filters in conditional point processes are conventionally fixed and thus do not directly model input-dependent spiking variability, though dependence on observed and unobserved covariates [91] and switching filters based on discrete states [23] have been considered. Recent work has moved away from parametric filters to nonparametric Gaussian processes [18], which can be extended to flexibly model dynamic filters as functions of external covariates with a spatio-temporal Gaussian process. However, any modulation of the filter will no longer permit fast convolutions, and such models will be computationally expensive as the filter needs to be recomputed every time step. The direct nonparametric estimation of conditional intensity functions based on maximum likelihood has been explored [13, 82], but scalable Bayesian approaches have remained absent.

### Contribution

To enable flexible modeling as well as modulation of instantaneous point process statistics for analyzing neural spike train variability, we introduce the Bayesian nonparametric non-renewal process (NPNR). NPNR builds on sparse variational Gaussian processes and defines a nonparametric prior over conditional interspike interval distributions, generalizing modulated renewal processes with nonparametric renewal densities and spike-history dependencies beyond renewal order. In particular, our point process can flexibly model modulations of not only spiking intensity but also variability. We validate our model using parametric inhomogeneous renewal processes, recovering conditional interspike interval distributions and identifying renewal order in spike-history dependence. On neural data from mouse thalamus and rat hippocampus, our method has competitive predictive power while being superior in capturing interspike interval statistics from the non-stationary data. In particular, our method provides instantaneous measures of spike train variability that are modulated by covariates, and shows rich variability patterns in both datasets consistent with previous studies at coarser timescales. We provide a JAX [4] implementation of our method as well as established baseline models within a scalable general variational inference scheme. ^1^

## 2 Background

We start with a brief overview of the theoretical foundations and related point process models, as well as their combination with Gaussian processes to introduce non-stationarity.

### 2.1 Temporal point processes

Statistical modeling of events that occur stochastically in time is handled by the general framework of temporal point processes [59, 72]. Denoting the number of events that occurred until time *t* by *N* (*t*), a temporal point process model is completely characterized by its conditional intensity function (CIF)

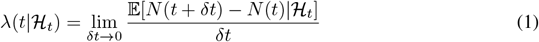

where *λ*(*t*)*δt* is the probability to emit a spike event in the infinitesimal interval [*t, t* + *δt*) conditioned on ℋ_*t*_, the spiking history before ℋ*t*. We can write the point process likelihood for a single neuron spike train consisting of an ordered sequence of *S* spike events at times *t*_*i*_ as [5]

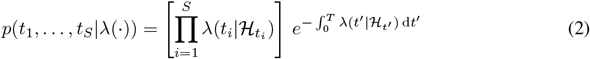

In neuroscience applications, one often wants to describe modulation of the point process statistics with some time-varying covariates ***x***(*t*), such as animal head direction or body position, which leads to a generalized CIF *λ*(*t*|ℋ_*t*_, ***x***_≤*t*_). Several classes of models have been proposed that are defined by particular restrictions on the functional form of *λ*(*t*|ℋ_*t*_, ***x***_≤*t*_).

#### 2.1.1 Inhomogeneous renewal processes

##### Renewal assumption

The statistical model in Eq. 2 describes dependencies between all spikes. One common simplification is the renewal assumption: interspike intervals (ISIs) ∆^(*i*)^ = *t*_*i*+1_ − *t*_*i*_ are drawn i.i.d. from an interval distribution called the renewal density *g*(∆; *θ*) with parameters *θ*. This induces a Markov structure *p*(*t*_*i*_|*t*_*i*−1_, *t*_*i*−2_, …) = *p*(*t*_*i*_|*t*_*i*−1_) in the spike train likelihood

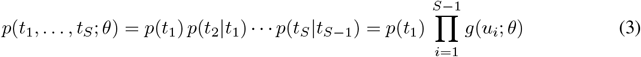

Common renewal densities used for neural data are the exponential (equivalent to a Poisson process), gamma, and inverse Gaussian distributions [2, 6].

##### Hazard function modulation

Non-stationary point processes need to model changes in statistics with time, and combined with the renewal assumption one obtains inhomogeneous renewal processes. A classical approach that dates back to Cox [14] is to modulate the hazard function (Appendix A)

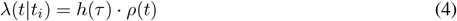

with time since last spike *τ* = *t*− *t*_*i*_ and the modulation factor *ρ*(*t*). In our context, this can be replaced by some function of covariates *ρ*(***x***_*t*_) [43]. A multiplicative interaction between *ρ* and *h* as above is typically considered, though this framework allows general parametric forms [71, 79].

##### Rate-rescaling

Another approach that has been widely applied in the neuroscience community is rate-rescaling [2, 6], closely related to time-rescaling (see Appendix B.3). Here, modulation is achieved with a rate function *r*(***x***_*t*_) ≥ 0 that transforms time *t* into rescaled time 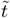

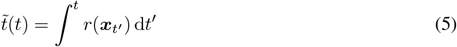

This maps all spike times *t*_*i*_ to rescaled times 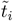, and will be one-to-one as long as *r*(*t*) *>* 0. By modeling the rescaled ISIs 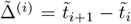 as drawn from a stationary renewal density *g*(·), we obtain an inhomogeneous renewal process from a homogeneous one. The CIF becomes dependent on the covariate path since last spike 𝒫_*k*_ = {***x***(*u*)|*u* ∈ (*t*_*i*_, *t*])}, see Appendix B.3.

#### 2.1.2 Conditional point processes

##### Conditional Poisson processes

The renewal assumption ignores correlations between ISIs, which generally are observed in both the peripheral and central nervous system [1, 25] and can be computationally relevant for signal detection and encoding [8, 9, 22, 69]. Going beyond Markovian dependencies, a tractable approach, similar to Hawkes processes [51], is to introduce a causal linear filter *h*(*t*) that is convolved with spikes and added to the log CIF, giving conditional Poisson processes

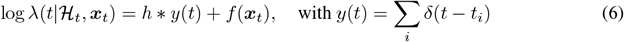

where ∗ denotes temporal convolution. These models are closely linked to mechanistic integrate-and-fire models [52, 85] and have a long history in the neuroscience literature [18, 31, 35, 47, 62, 66, 83], appearing as generalized linear models (GLMs) and spike response models (SRMs).

##### Conditional renewal processes

An even more expressive model can be obtained by replacing the Poisson spiking process with a rate-rescaled renewal process. This results in a conditional renewal process, where the rate function has history dependence *r*(*t*|H_*t*_, ***x***_*t*_) as in Eq. 6.

### 2.2 Gaussian process modulated point processes

Gaussian processes (GPs) represent a data-efficient alternative to neural networks, which have been widely used to model the CIF [59, 72, 90]. When combining GPs with point process likelihoods, the resulting generative model leads to doubly stochastic processes for event data. Placing a Gaussian process prior [87] over the log intensity function leads to the classic log Cox Gaussian processes [14, 55], and in the same spirit one can modulate renewal hazard functions [79] or perform rate-rescaling [16] with GPs. Such constructions form the basis of many widely used Bayesian neural encoding models for spike trains, both for modeling single neuron responses [15, 16, 68] as well as population activity [19, 93]. Combining the flexibility offered by renewal and conditional point processes with GP rate or modulation functions within a variational framework has been impeded by the fact that the original papers were built on a GLM framework with parametric covariate mappings [6, 7, 18, 65]. To provide a fair comparison of NPNR to these baselines, we implement a general scalable variational inference framework for the construction and application of such models (see Appendix B for details on the baseline models).

## 3 Method

We now introduce the nonparametric non-renewal (NPNR) process and present the approximate Bayesian inference scheme used for model fitting, noting connections to related works in the literature. Our NPNR model provides a nonparametric generalization of modulated renewal processes beyond renewal order, and adds suitable inductive biases for neural spike train data. It implicitly defines a flexible prior over conditional ISI distributions that can be computed using pathwise conditioning, which enables one to analyze spiking variability modulation with minimal parametric constraints. Furthermore, the Bayesian framework provides an elegant data-driven approach to inferring the lagging ISI order of the spike-history dependence.

### 3.1 Generative model

#### Conditional intensity surface priors

To obtain flexible modulated point process models, we directly model the CIF, or more precisely its logarithm, of the form

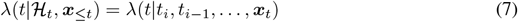

where *t*_*i*_ is the most recent spike at current time *t*. First considering the renewal case, we note the spatio-temporal structure in the log CIF using time since last spike *τ* = *t* − *t*_*i*_ is

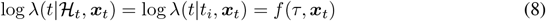

which suggests placing a spatio-temporal GP prior on the log CIF to describe a log intensity surface

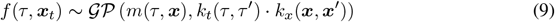

This generalizes the parametric forms of modulation considered in previous approaches [43, 79], in particular allowing modulation of the effective instantaneous renewal density by covariates ***x***_*t*_. We can introduce lagging ISIs covariates ∆_*k*_(*t*) with lag *k* as depicted in Fig. 1A to extend the model to a non-renewal process

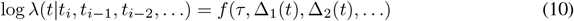

and for a maximum ISI lag *K* we denote the lagging ISIs **∆**_*t*_ = [∆_1_(*t*), …, ∆_*K*_(*t*)] to obtain

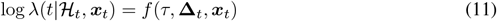

**Figure 1:**
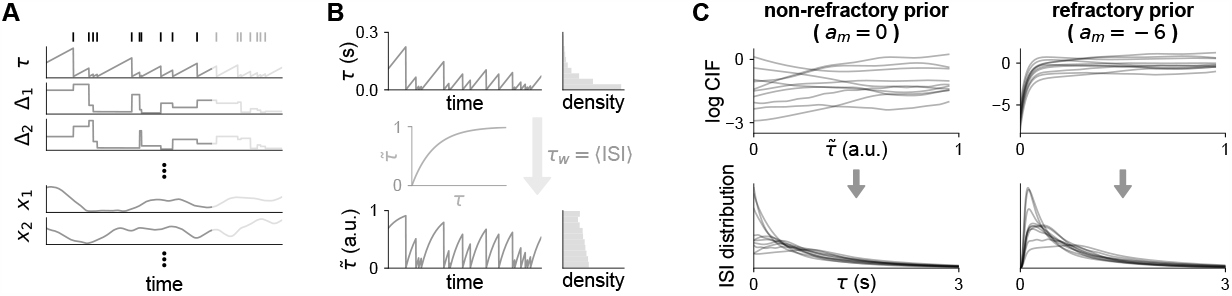
Schematic of our proposed model. **(A)** Time since last spike *τ* and lagging ISIs **∆** for an observed spike train (top row) alongside covariates ***x*. (B)** Illustration of the time warping procedure. We fix the warping parameter *τ*_*w*_ to the empirical mean ISI, which leads to more uniform distributions 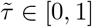 suitable for a stationary GP kernel. **(C)** Prior samples from the generative model for two values of *a*_*m*_ characterized by the lack and presence of a refractory period. The transformation Eq. 15 links the log CIF (top rows) with conditional ISI distributions (bottom rows).

#### Inductive biases for neural data

For neural spiking data, there are biological properties to consider for building a more realistic prior. Firstly, neurons have refractory periods immediately following a spike, though in practice neural recordings may not respect this due to contamination in spike sorting [38]. Another potentially useful inductive bias is that changes in the spiking intensity of neurons fluctuate mostly at shorter ISI timescales [32], whereas at longer delays they tend to be temporally smoother. The latter suggests non-stationary GP kernels to be more suitable for modeling the spike-history dependencies [18]. However, non-stationary kernels do not allow straight-forward use of random Fourier features for evaluating GP posterior function samples at many locations with pathwise conditioning [88, 89]. This in particular is needed to compute the conditional ISI distributions *g*(*τ*| …) in Eq. 15, see Appendix B.5 for details. To achieve the desired non-stationarity for modeling Eq. 11 while maintaining the ability to draw samples using pathwise conditioning, we apply time warping on *τ* from [0, ∞) → [0, 1] with some warping timescale *τ*_*w*_

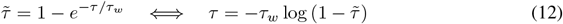

and place a stationary Gaussian process prior over the warped temporal dimension 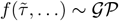 with a temporal kernel 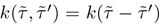. This transformation is monotonic (see Fig. 1B), and hence we can easily compute the transformation on the CIF

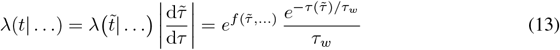

Similarly, we apply time warping to the ∆_*k*_ dimensions on which we also place stationary kernels 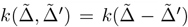. We note that unlike spike-history filters in conditional point processes (Eq. 6) which do not change with inputs, the resulting coupling to past activity in Eq. 11 is dependent on covariates ***x***_*t*_. This allows one to capture spiking variability modulation via the conditional ISI distribution perspective discussed below. The refractory nature of real neurons can be addressed by the mean function

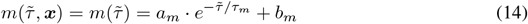

with parameters *a*_*m*_, *τ*_*m*_, *b*_*m*_, where refractory periods can be modeled with large negative *a*_*m*_.

#### Conditional ISI distributions

Instead of looking at the CIF, we can view the model as a prior over conditional ISI distributions as depicted in Fig. 1C using the relation (see Appendix A)

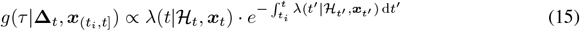

where one drops the dependence on ℋ_*t*_ in *g* for the modulated renewal case. If one fixes the lagging ISIs and picks a constant covariate path *g*(*τ* |**∆**_∗_, ***x***_∗_), this can be interpreted as an instantaneous ISI distribution of a neuron at the conditioned inputs **∆**_∗_ and ***x***_∗_. Moments of the conditional ISI distribution are computed using Gauss-Legendre quadratures in warped time (Appendix B.5)

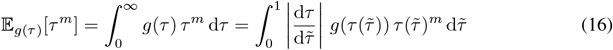

and this can be used to compute tuning curves of spike train statistics, such as the mean ISI 𝔼 [*τ*] and coefficient of variation 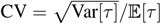 [58], as a function of ***x***_∗_. This approach generalizes the homogeneous case considered in the literature, and in particular allows one to compute instantaneous measures of non-stationary spike train variability that are otherwise non-trivial to estimate [58, 75].

### 3.2 Inference

#### Temporal discretization

The generative model is formulated as a continuous time model. In practice, neural and behavioural data are typically recorded with finite temporal resolution at small regular intervals ∆*t*. The cumulative intensity integral has to be approximated by a sum, though note that directly modeling the cumulative hazard function [59] elegantly avoids this for purely temporal point processes. Spike times are now discretized as a binary vector ***y*** = [*y*_1_, …, *y*_*T*_] where *y*_*t*_ = 1 if *t* has a spike event, zero otherwise. Overall, this discretizes the point process likelihood Eq. 2 as

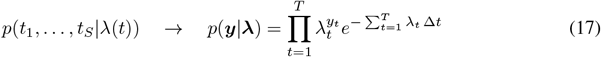

where we have *T* time steps in total. Note that the discretization scheme implies we do not have observations at *τ* = 0, since the time step immediately after an observed spike has *τ* = ∆*t*.

#### Variational lower bound

We use stochastic variational inference [39] with batches obtained from consecutive temporal segments and sparse variational GPs [37], giving the loss objective

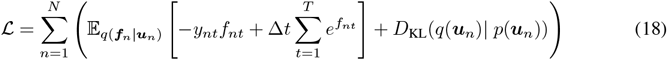

where *n* indexes neurons (of which there are *N*)^2^, ***u*** = [*u*_1_, …, *u*_*M*_] denotes the set of *M* inducing points, *p*(***u***) the GP prior at inducing locations, *q*(***u***) the variational posterior, and *q*(***f u***) the conditional posterior (see Appendix B.1 for details). Combined with temporal mini-batching to fit batch segments of length *T*, we can fit to very long time series given the 𝒪 (*N T M* ^2^ + *N M* ^3^) computational complexity. We also no longer rely on computing hazard functions of parametric renewal densities to obtain the CIF, which can be numerically unstable. Modulated renewal processes instead rely on a specialized thinning procedure [79], but we take a more scalable and general variational approach. Overall, we optimize the kernel hyperparameters, variational posterior mean and covariance, inducing point locations, and mean function parameters *a*_*m*_, *b*_*m*_ and *τ*_*m*_ using gradient descent with Adam [46] (see Appendix C for details). The time warping parameter *τ*_*w*_ is fixed in our experiments to the empirical mean ISI, and the hyperparameter *K* is fixed and chosen in advance (see also subsection on automatic relevance determination below).

#### Automatic relevance determination

The Bayesian framework with GPs enables us to perform automatic relevance determination (ARD) over the input dimensions to automatically select relevant input [37, 80]. Applied to lagging ISI dimensions in our NPNR model, this provides an elegant approach to making a data-driven renewal assumption and generally determining the spike-history dependence of the CIF. We choose to fix *τ*_*w*_ to the empirical mean ISI as shown in Fig. 1B (rather than learning it) to achieve interpretability of kernel timescales for ARD (Fig. 6A) at a small cost of performance (Fig. 12). For a chosen maximum lag *K*, there is no need for manual selection of the history interaction window as for GLM spike-history filters, though recent work on nonparametric GLM filters provides a related window size selection procedure [18]. In the spirit of Bayesian models, we choose *K* to give a sufficiently high capacity model [40, 80] to be able to flexibly capture history dependence, as seen in panel D of Fig. 3 and Fig. 4.

**Figure 2:**
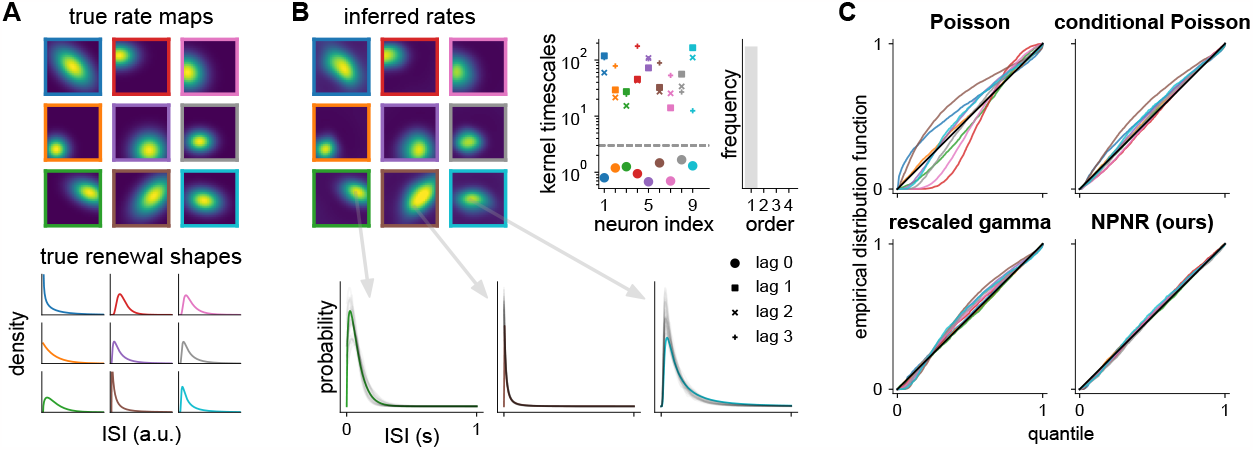
Validation on synthetic data. **(A)** True rate maps (brighter is higher) defined over a square environment (top) and base renewal densities (bottom) in each column given by gamma (left), log normal (middle) and inverse Gaussian (right) distributions with various shape parameters. Each color corresponds to a separate neuron. **(B)** Posterior mean rate maps (top left) and conditional ISI distribution samples in gray overlaid on true renewal densities at various locations (bottom) for the NPNR process fit to synthetic data. The relevance boundary (dotted line) for kernel timescales (top right) is placed at *l* = 3 (dimensionless). **(C)** QQ-plots of fitted models (each curve is a neuron).

**Figure 3:**
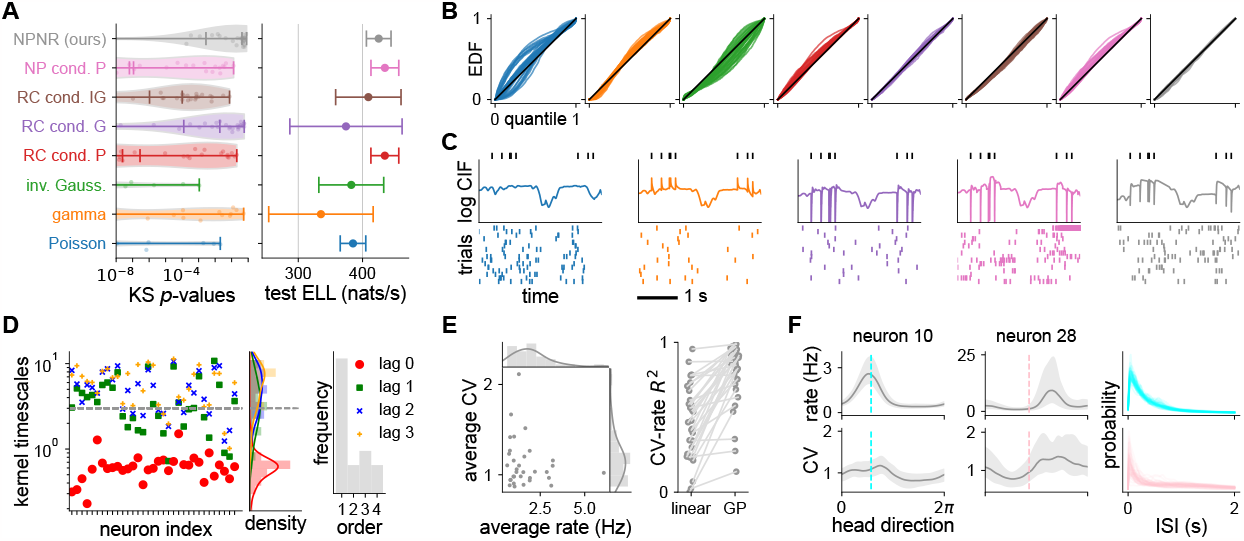
Application to mouse head direction cell data. **(A)** Violin plot of KS *p*-values per neuron (left, lines marking quartiles) and test expected log likelihoods with errorbars showing s.e.m. across test folds (right). Larger values in both metrics indicate better model fit to data. **(B)** QQ-plots for various models (each curve is a neuron) identified by color (panel A left). **(C)** Predicted log CIF (middle) for an observed spike train (top) and posterior spike train samples (bottom) conditioned on the same covariates ***x***_*t*_ for various models identified by color. **(D)** *Left:* temporal kernel timescales for *τ* (lag 0) and ∆_*k*_ dimensions with the relevance boundary at *l* = 3 (dimensionless). *Right:* histogram of “ISI-order” (1 + largest lag *k* for which *k* is below the boundary) across neurons. **(E)** Time average of estimated instantaneous rates and CVs from the training data (left) and *R*^2^ values of CV-rate regression with a linear and a GP model (right). **(F)** Posterior median and 95% intervals of tuning curves over head direction for the rate and CV, with posterior ISI distribution samples (right) at dashed locations.

**Figure 4:**
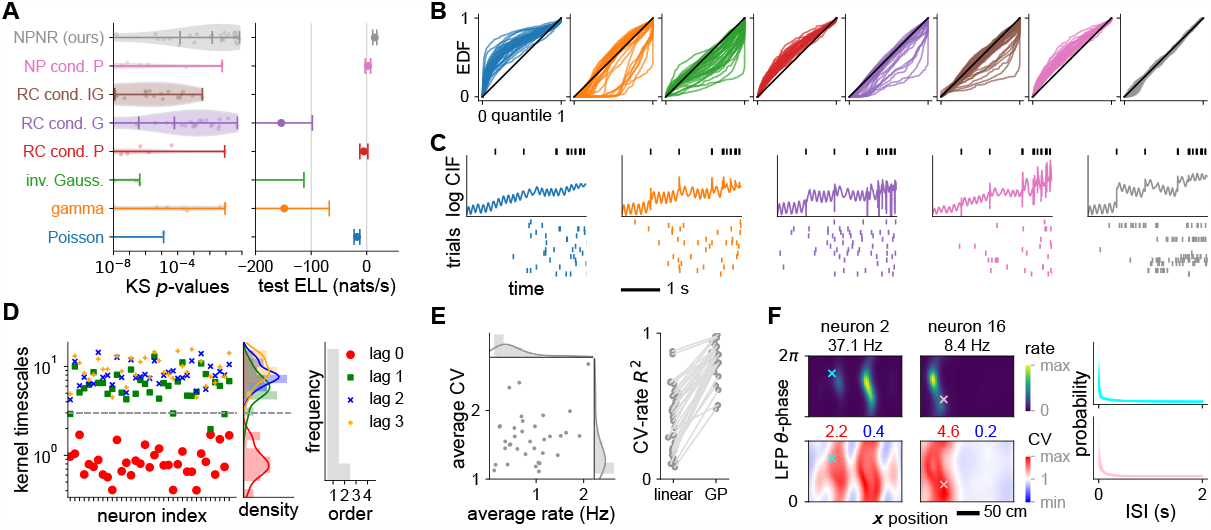
Application to rat hippocampal place cell data. **(A)**-**(E)** Similar to Fig. 3A-E. **(F)** Posterior mean tuning maps over *x*-position and *θ*-phase for the rate and CV (left) for left-to-right runs (based on head direction) with posterior ISI distribution samples (right) at marked locations.

**Figure 5:**
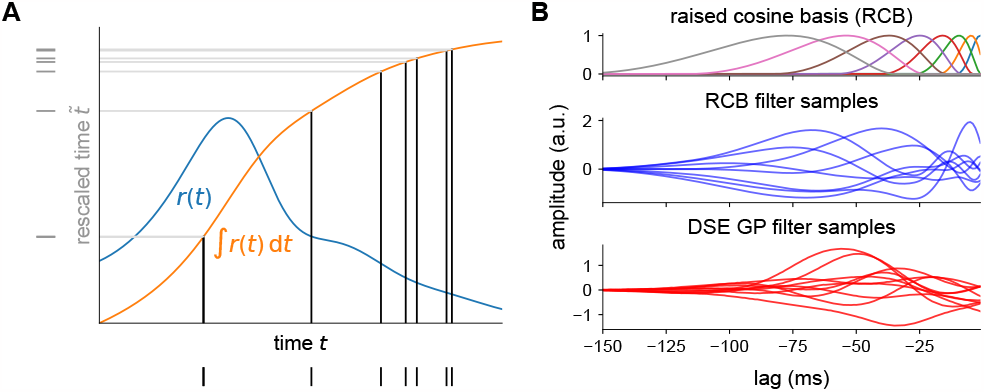
Details on baseline models. **(A)** Rate-rescaling procedure. The cumulative integral (orange) of a non-negative rate function (blue) define the monotonic mapping between real time *t* and warped time 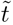. **(B)** Spike-history filters in conditional point processes. The raised cosine basis (top) is used to construct filters describing interactions with past spikes (middle). Nonparametric GP filters using the DSE kernel encode a similar temporal structure without explicit parameterization (bottom).

**Figure 6:**
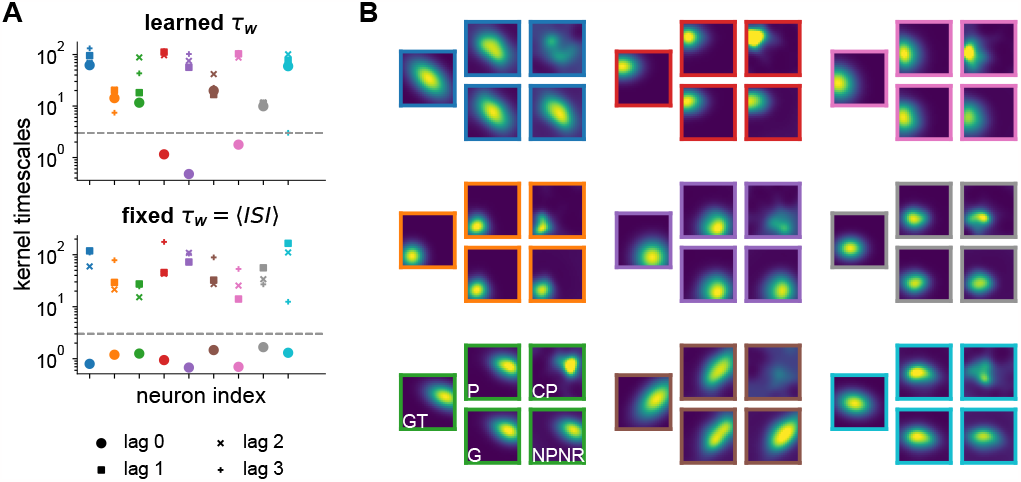
Additional results from synthetic experiments. **(A)** Comparing history-dependence ARD for fixed and learned time warping parameter *τ*_*w*_. The interference between *τ*_*w*_ and the kernel timescales in our model causes failure of ARD when learning *τ*_*w*_, as kernel timescales are not longer identifiable with effective temporal fluctuations in the log CIF. **(B)** Estimated rat maps from fitted models. Each subpanel shows rate maps (brighter is higher) for one synthetic neuron, with the ground truth (GT) on the left and grouped on the right inferred maps from the Poisson (P), conditional Poisson with raised cosine filters (CP), rate-rescaled Gamma (G) and the NPNR model.

## 4 Results

All datasets discretize spike trains and input time series at regular intervals of ∆*t* = 1 ms. We use a product kernel for *k*(***x, x***^*′*^) with periodic kernels for angular dimensions, and squared exponential kernels in other cases. For 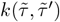 and 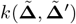, we pick a product kernel with Matérn-^3^*/*_2_ (see Fig. 12 for different kernel choices) and set the maximum ISI lag *K* = 3. For illustration, conditional ISI distributions and corresponding tuning curves are computed by fixing **∆**_*k*_ to be the mean ISI per neuron. Firing rates are defined as 1*/* 𝔼 [*τ*], since this corresponds to the number of spikes fired per unit time in infinitely large time bins for a renewal process (Eq. 26). GP inducing points were randomly initialized, and for a fair comparison, all models used 8 inducing points for each covariate dimension (including temporal dimensions *τ* and **∆** in the NPNR process). For each experiment, we repeat model fitting with 3 different random seeds and pick the model with the best training likelihood. Further details on experiments are presented in Appendix C.

### 4.1 Validation on synthetic data

For validating our approach, we generate 1000 s of data using rate-rescaling [65] mimicking a place cell population of 9 neurons for an animal moving in a 2D square arena, each with a unique rate map and renewal density (Fig. 2A, details in Appendix C). The models applied are baseline Poisson, raised cosine filter conditional Poisson and rate-rescaled gamma processes (Appendix C), and our NPNR process. Note that the rescaled gamma process is within-model class for 3 of the synthetic neurons. Inferred conditional ISI distributions and rate maps of our NPNR process in Fig. 2B are close to ground truth, showing the ability of our model to capture modulated spiking statistics drawn from various parametric families. To assess how well ISI statistics are captured, we apply time-rescaling using the GP posterior mean functions (Appendix A) which we visualize with quantile-quantile (QQ) plots [6] in Fig. 2C. Again, we see an excellent fit of our model compared to baseline models, indicating that only the NPNR is capable of satisfactorily capturing the empirical ISI statistics. Learned temporal kernel timescales of the NPNR process in Fig. 2B show a clear separation between the time since last spike *τ* dimension (lag 0) and lagging ISI **∆** dimensions (lag ≥ 1) with the dotted relevance boundary at *l* = 3 (dimensionless), as expected for renewal processes.

### 4.2 Neural data

Now we apply our method to head direction cells in freely moving mice [63, 64] and place cells in rats running along a linear track [54]. We select 33 units from the mouse and 35 units from the rat data, which leads to around 36 and 68 million data points to fit in the training set, respectively (see Appendix C for preprocessing details and Appendix B.2 on data scaling). Experiments involve fitting to the first half of a dataset (∼18 min. for mouse, ∼32 min. for rat), and testing on the second half split into 5 consecutive segments. The split into 5 test folds is used to quantify dataset variability. For prediction, we evaluate the expected log likelihood summed over all neurons

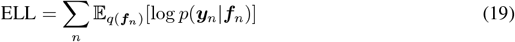

using Gauss-Hermite quadrature with 50 points (Monte Carlo for renewal processes, see Appendix B). To assess goodness-of-fit to ISI statistics, we compute QQ plots as before and apply the Kolmogorov-Smirnov (KS) test, giving a *p*-value per neuron that indicates how likely the data came from the model (Appendix A). Baselines are the inhomogeneous Poisson (P), rate-rescaled gamma (G) and inverse Gaussian (IG) renewal [6, 83], raised cosine (RC) filter conditional Poisson [66, 85] and renewal [65], and nonparametric (NP) filter conditional Poisson processes [18] (details in Appendix C).

#### 4.2.1 Mouse head direction cell data

We choose the animal head direction as our 1D input covariate *x*. From the KS test *p*-value distribution, we see that our model outperforms all baselines in capturing ISI statistics (Fig. 3A left, higher is better; Fig. 3B, QQ plots closer to diagonal). It performs competitively to conditional Poisson processes in terms of predictive performance (Fig. 3A right), but those models fail to capture ISI statistics. In addition, we note the spiking saturation in some samples of the nonparametric conditional Poisson model (Fig. 3C, purple) due to a known instability [3, 29]. Samples from our model (Fig. 3C, gray) exhibit visually similar spike patterns to the real spike train segment. Furthermore, kernel timescales in Fig. 3D show a sizable fraction of the population is characterized by a non-renewal spiking process.

##### Neural dispersion regimes

From Fig. 3E and F, we observe both under- and overdispersion (CV smaller and bigger than one) consistent with a previous study based on spike counts [48]. Estimated instantaneous rates and CVs in Fig. 3E are computed using the conditional ISI distribution evaluated along the time series of covariates in the training data. One can regress instantaneous CV against rate, and the CV-rate *R*^2^ shows some cells with near linear relations and some with nonlinear trends that can be captured by a GP. Despite that, many cells still show a low overall *R*^2^, implying there is no parametric relation. Most neurons increase CV with rate, while some show a slight linear decrease reminiscent of refractory Poisson processes (Eq. 31, see Fig. 7 for CV-rate patterns).

**Figure 7:**
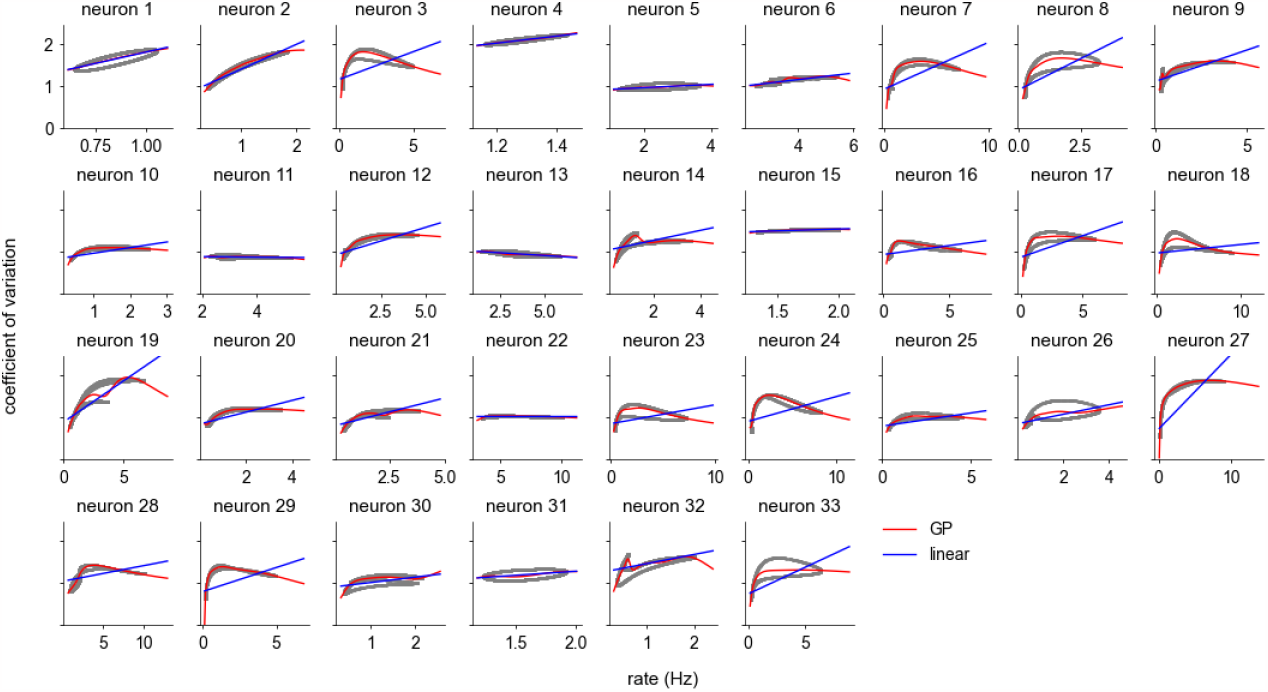
Spiking variability of head direction cells. Dots represent estimated instantaneous statistics computed using the posterior conditional ISI distributions. Linear and GP regression fits are shown overlaid on top of the dots. Note the one-dimensional trajectory that is traced out is due to the fact that we have 1D covariates ***x***_*t*_ (head direction).

#### 4.2.2 Rat place cell data

In this case, ***x*** is 3D consisting of the body position along the track, head direction, and local field potential (LFP) *θ*-phase. Our model significantly outperforms all baselines, having both a better KS test *p*-value distribution (Fig. 4A left, higher is better; Fig. 4B, QQ plots closer to diagonal) and predictive performance (Fig. 4A, right). Note that the ELLs for this dataset differ from those shown in Fig. 3A due to the fundamentally different (less predictable) spiking statistics of place cells compared to head direction cells (e.g. due to theta oscillations). As the nonparametric conditional Poisson model introduces nonparametric but *covariate-independent* variability patterns, these results highlights the importance of modeling *covariate-dependent* spiking variability. Note that the rate-rescaled renewal processes struggle to fit this data, with test ELLs of inverse Gaussian models below -200 nats/s (Fig. 4A right). Samples from our model (Fig. 4C, gray) show it captures the characteristic bursting nature of the real spike train segment. Kernel timescales in Fig. 4D show most cells are described well by a renewal process, different to mouse data Fig. 4D.

##### Capturing overdispersion

We see CV values in Fig. 4E higher than the mouse thalamus dataset, consistent with overdispersion of place cell discharge in 2D open field navigation [26]. Similar to the mouse data, the CV-rate *R*^2^ again shows there is generally no parametric relation. We also tend to observe larger increases in CV with firing rate compared to mouse data (Fig. 9).

**Figure 8:**
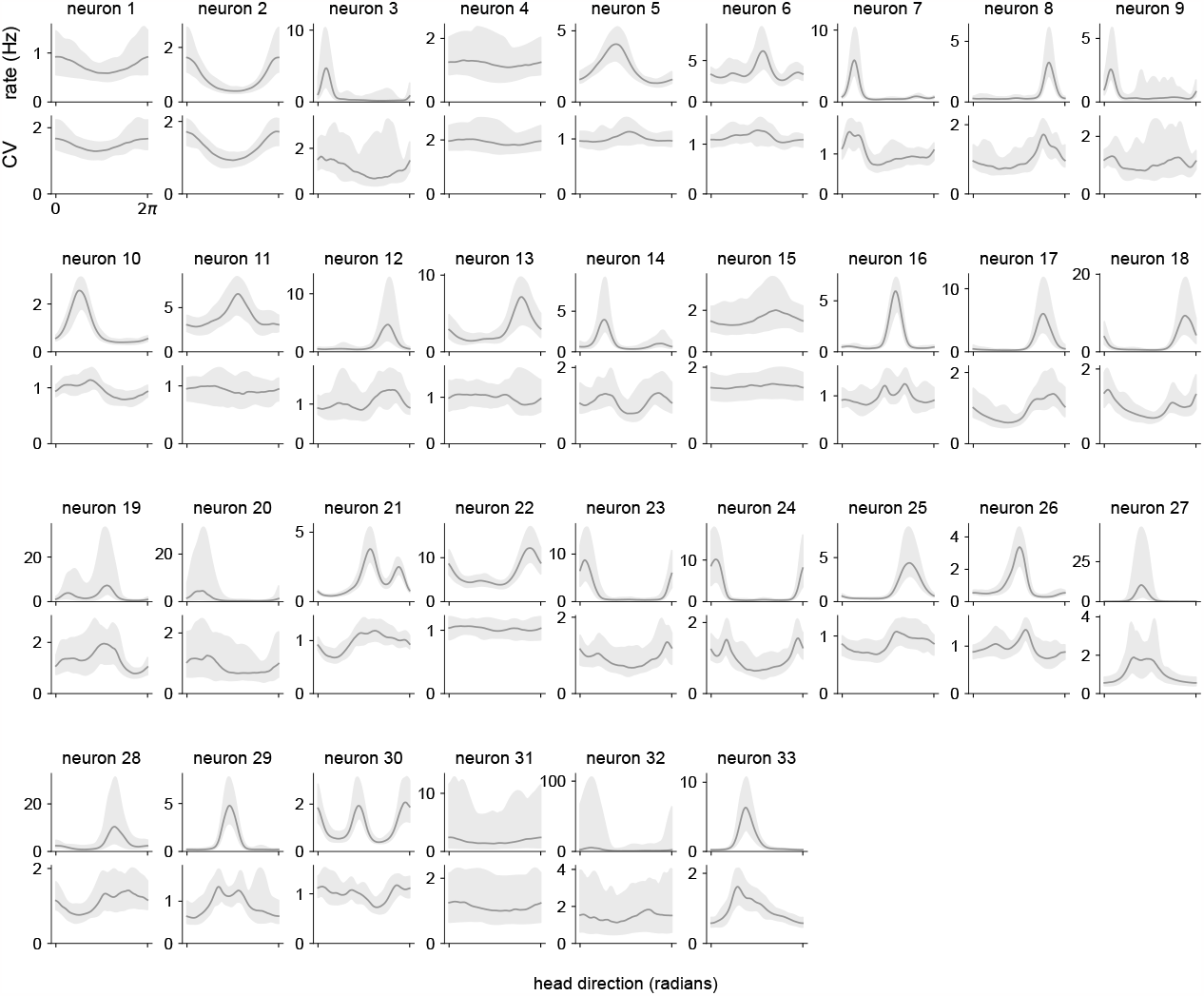
Tuning curves of head direction cells. Spike train statistics are computed from conditional ISI distribution samples. Lines show posterior medians, and shaded areas show 95% credible intervals.

**Figure 9:**
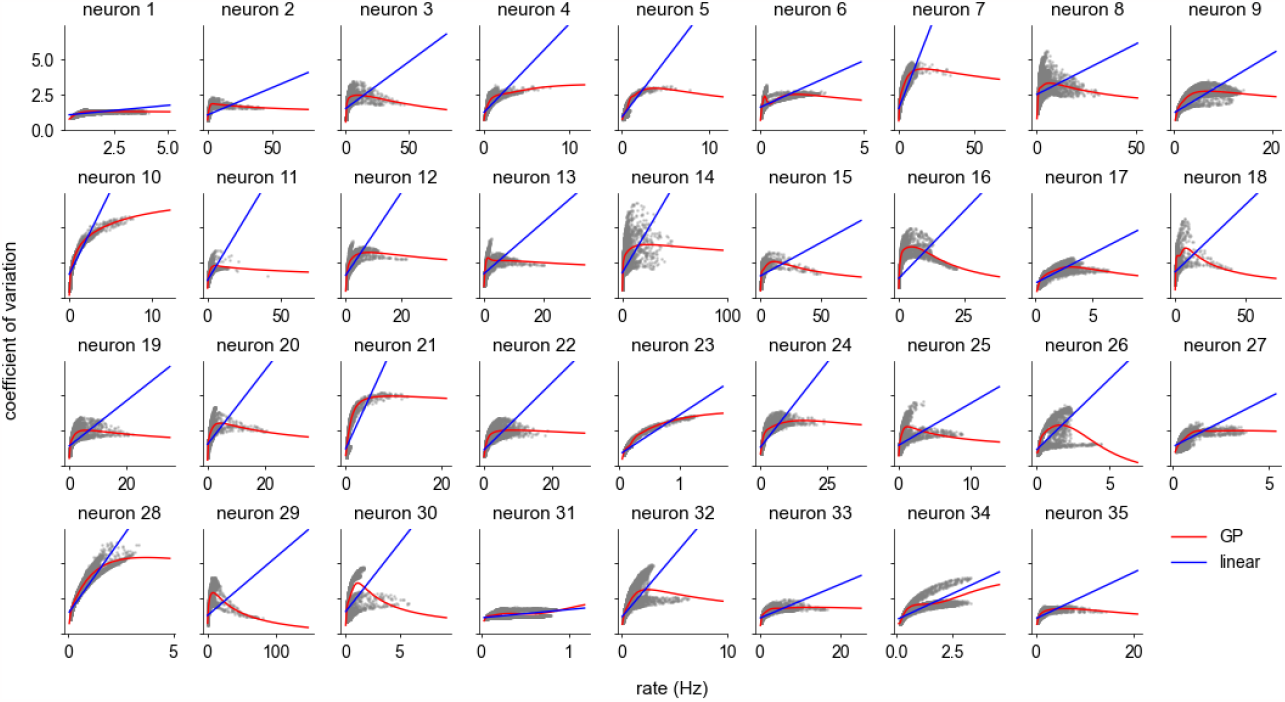
Spiking variability of place cells. Dots represent estimated instantaneous statistics computed using the posterior conditional ISI distributions. Linear and GP regression fits are shown overlaid on top of the dots.

##### *θ*-modulation and phase precession

Spiking activity modulation during *θ*-cycles [53] is prominent in rat hippocampus, and is visible here in the log CIF (Fig. 4C). We also see phase precession [76] in Fig. 4F (top), a classical example where spike timing relative to some rhythm has coding significance [33]. Our model enables one to extract not only spiking intensity but also variability, and shows that variability generally inherits the phase precession pattern (Fig. 4F bottom).

## 5 Discussion

### 5.1 Limitations and further work

#### ISI statistics

Apart from the coefficient of variation, there are other ISI statistics that characterize spiking dynamics aspects such as bursting or regularity. Of particular interest is the local coefficient of variation [75], which involves joint statistics of consecutive ISIs (∆^(*i*)^, ∆^(*i*−1)^) that can be computed from our model (see Appendix D). The same applies to serial correlations [25], which may provide insights into biophysical details [77]. Furthermore, quantifying the shape of ISI distributions is of interest as it is associated with various properties of the underlying neural circuit dynamics [61].

#### Neural correlations

To capture correlations in multivariate spike train data, direct spike couplings as in GLMs are less suitable for current neural recordings compared to latent variable models due to the sparse sampling of populations by electrodes [50]. Combining the latter alongside observed covariates [48] with our point process provides a powerful framework for capturing correlations [81], which can have significant impact on neural coding [56]. To perform goodness-of-fit tests, the Kolmogorov-Smirnov test with time-rescaling can be extended to the multivariate case [30, 92].

### 5.2 Conclusion and impact

We introduced the Bayesian nonparametric non-renewal (NPNR) process for flexible modeling of variability in neural spike train data. On synthetic renewal process data, NPNR successfully captures spiking statistics and their modulation by covariates, and finds renewal order in the spike-history dependence. When applied to mouse head direction cells and rat hippocampal place cells, NPNR has competitive or improved predictive performance to established baseline models, and is superior in terms of capturing ISI statistics, establishing the importance of capturing covariate-dependent variability. NPNR-based analyses recover known behavioral tuning, while also revealing novel patterns of spiking variability at millisecond timescales that are compatible with count-based studies.

Neural firing rates traditionally characterize most computational functions and information encoded by neurons [16, 17, 33], but recent work on V1 [20, 27, 36, 60] and hippocampal place cells [84] have started to assign computationally well-defined roles to variability in the context of representing uncertainty. Our method introduced in this paper is a principled tool for empirically characterizing neural spiking variability and its modulation at the timescales of individual spikes, and we hope our model will be useful for revealing new aspects of neural coding. Such findings are foundational to advances in computational and theoretical neuroscience, and may have downstream practical applications in designing and improving algorithms for brain-machine interfaces.

## Acknowledgments and Disclosure of Funding

This work was supported by the Cambridge European and Wolfson College Scholarship by the Cambridge Trust (D.L.) and by the Wellcome Trust (Investigator Award in Science 212262/Z/18/Z to M.L.). We are grateful to Kristopher Jensen, Marine Schimel and Valentina Njaradi for helpful comments on the manuscript. We would also like to thank Alexander Terenin and Jonathan So for helpful discussions.

## Supplementary Material

### A Point process theory

#### A.1 Point process definitions

From the conditional intensity function (CIF) defined in Eq. 1, we can obtain the survival function *S*(*τ*) based on the flux of probability using time since last spike *τ* = *t* − *t*_*i*_

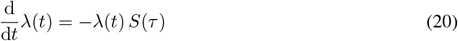

which quantifies what the probability is to have no events from *t*_*i*_ up to time *t*. For the relation between the interspike interval (ISI) distribution *g*(*τ*), we require an event to occur within the infinitesimal window [*t, t* + *δt*) which is proportional to multiplying the CIF with the survival function [14]

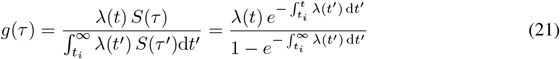

Another point process quantity that appears in the literature is the hazard function [9, 31], which is related to the CIF by number at risk. In our case of modeling neural spike trains, this is the same as the CIF. The hazard function is often considered in renewal processes, where we have the relation

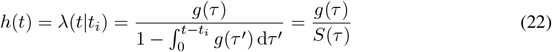

#### A.2 Time-rescaling and Kolmogorov-Smirnov goodness-of-fit test

##### A.2.1 Time-rescaling

Time-rescaling is analogous to Eq. 5 but uses the CIF instead of a rate function

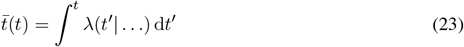

with rescaled time 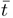. The rescaled ISIs should then be exponentially distributed if the ISIs are computed from spike train samples of the point process with the given CIF [27].

##### A.2.2 Quantile-quantile plots and dispersion

The time-rescaled ISIs 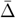 can be transformed with the cumulative density function of the unit exponential distribution into quantiles in the range [0, 1]

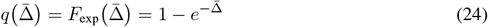

which should be uniformly distributed if we are working with samples from the point process with the given CIF. One can visualize the empirical distribution of quantiles from rescaled ISIs using a quantile-quantile (QQ) plot as in Fig. 2C. There we plot the empirical distribution function *F* (*q*) of quantiles *q*. If the point process model matches the empirical ISIs well, the QQ plot curve will follow the diagonal (i.e. the empirical distribution function of a uniform random variables).

The terms over- or underdispersion describe empirical quantile distributions that do not match the point process model. Overdispersion is associated with too many extreme quantile values near 0 or 1. On the other hand, underdispersion involves an excess of quantiles close to the median activity *q* = ½. These two forms of dispersion will show up characteristically on QQ plots, with underdispersion associated to an S-like curve and overdispersion its reflection along the diagonal. In neuroscience, the reference point process is conventionally taken to be an intensity-matched Poisson process. Over- or underdispersion then refers to to spiking activity that is more irregular or regular temporally than a Poisson process, respectively.

##### A.2.3 Kolmogorov-Smirnov test

The intuition from QQ plots can be formalized using a statistical test called the Kolmogorov-Smirnov goodness-of-fit test. The test statistic is defined by

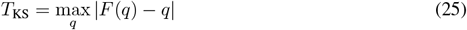

which represents the biggest vertical distance of the empirical distribution function to the diagonal. Since we deal with a different number of ISIs for each neuron, we report *p*-values of the observed *T*_KS_ [30] in Fig. 3A and Fig. 4A to have a comparable measure across neurons. Low *p*-values indicate the null hypothesis (the proposed point process model) should be rejected, or equivalently does not match the empirical data well. In the literature, this is a standard test for assessing goodness-of-fit of neural spike train models [8, 23].

#### A.3 Renewal processes

##### A.3.1 Firing rates and ISIs

The law of large numbers for renewal processes [24] shows that for a Markov renewal process

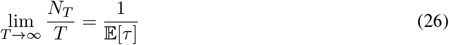

where *N*_*T*_ is the number of events in the interval [0, *T*]. This relation can be interpreted as the asymptotic firing rate being equal to the reciprocal mean ISI. Note for finite bin sizes *T*, this generally does not hold and the two quantities will differ depending on the ISI distribution shape.

##### A.3.2 Parametric renewal density families

Below we give the parametric densities for renewal processes used in the paper. Note that for model fitting, we rescale the density such that we obtain unit mean by dividing *τ* with 𝔼 [*τ*].

###### Gamma

The Gamma renewal process is obtained from picking a Gamma renewal density

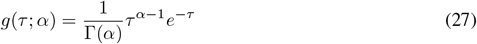

which is parameterized by the shape *α*. Note the Poisson process corresponds to *α* = 1 i.e. the exponential distribution. This has mean 𝔼 [*τ*] = *α*. The shape parameter *α* acts as a “tuning knob” for spiking randomness, with *α <* 1 leading to overdispersed and *α >* 1 underdispersed activity.

###### Inverse Gaussian

The inverse Gaussian is ased on the

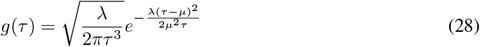

where the parameter *λ* is absorbed by the rate-rescaling transform. Hence we take *λ* = 1 without loss of generality for neuroscience applications. This has mean 𝔼 [*τ*] = *μ*.

###### Log-normal

The log-normal distribution is named as the logarithm of the random variable is normally distributed

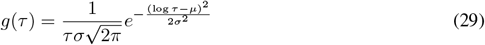

noting that the rate-rescaling absorbs the parameter *μ*. Hence in neuroscience applications, the renewal density will be parameterized by *σ* with *μ* = 0. This has mean 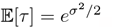.

###### Refractory Poisson

The Poisson process is commonly used, but it does not account for the refractory nature of real neurons. A simple modification is to introduce an absolute refractory period

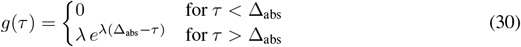

which introduces a dependence on the previous spike time, giving a renewal process with

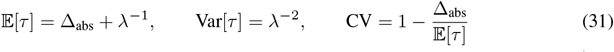

Note the linear relation between the coefficient of variation CV and the inverse mean ISI 𝔼 [*τ*]^−1^.

##### A.3.3 Hazard functions and asymptotic limits

To evaluate the CIF for renewal processes, we need to compute the hazard function as discussed above. Numerically, this can be challenging as we need to compute the fraction of the renewal density and the survival function (Eq. 22) which both tend to 0 for large *τ*, and hence we compute a truncation of the asymptotic series instead. Note that the survival function for renewal processes *S*(*τ*) = 1 − *C*(*τ*) with *C*(*τ*) cumulative density function of *g*(*τ*).

###### Gamma

The cumulative density function is

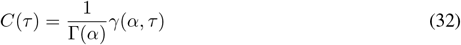

where *γ*(·, ·) denotes the lower incomplete Gamma function. The survival function satisfies

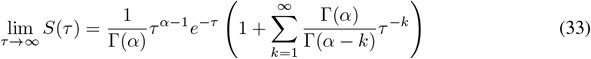

and the hazard function becomes

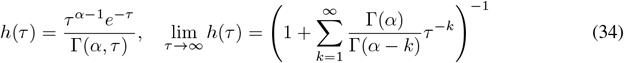

using the upper incomplete Γ(*α, x*) = 1 − *γ*(*α, x*)

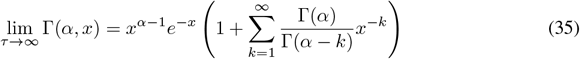

###### Inverse Gaussian

The cumulative density function is

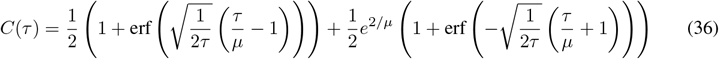

and thus

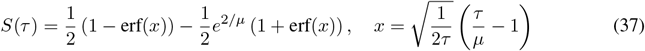

and the survival function has asymptotic limit

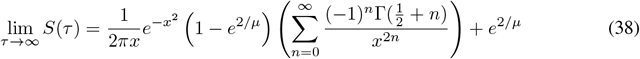

giving the hazard function limit for *τ* → ∞

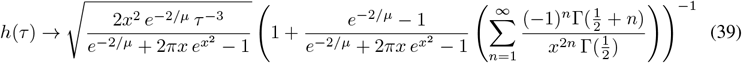

###### Log normal

The cumulative density function is

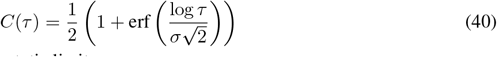

and the survival function has asymptotic limit

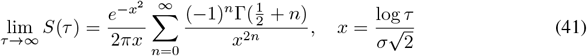

giving the hazard function

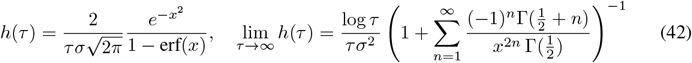

using 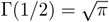 and the error function expansion

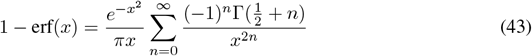

### B Implementation details

#### B.1 Sparse variational Gaussian processes

##### B.1.1 Gaussian processes as priors over functions

Gaussian processes (GPs) are a class of widely used Bayesian nonparametric models for modeling unknown functions [36]. Briefly, a Gaussian process 𝒢 𝒫 (*m*(·), *k*(·, ·)) is defined by a mean and covariance function *m*(·) and *k*(·, ·), and specifies a prior over functions *f* (·) ∼ 𝒢𝒫 such that for any set of input locations 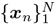, the function values ***f*** = [*f* (***x***_1_), …, *f* (***x***_*N*_)] satisfy

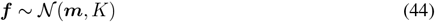

with mean vector ***m*** = [*m*(***x***_1_), …, *m*(***x***_*N*_)] and covariance matrix *K*_*ij*_ = *k*(***x***_*i*_, ***x***_*j*_).

##### B.1.2 Sparse approximation

GPs suffer from an *O*(*N* ^3^) computational bottleneck for inference [36], where *N* is the number of data points. In addition, closed-form inference and prediction are not possible for non-Gaussian likelihoods as used in this paper. To approximate intractable Gaussian process posteriors in non-conjugate settings, as well as remove the *O*(*N* ^3^) bottleneck for inference and posterior evaluation, we use variational inference along with a sparse approximation of the variational psoterior. The latter refers to parameterizing an approximate posterior as a conditional Gaussian distribution, conditioned on *M* additional function points at locations 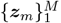 called inducing points. This leads to a sparse posterior in the sense that *M < N*, while the generative model is augmented given by the joint distribution (where we set the mean *m*(·) = 0 for convenience without loss of generality)

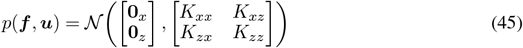

Following [33], we directly parameterize the posterior over the function values at inducing points

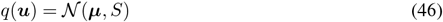

giving a joint posterior for the GP model

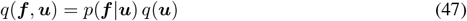

The resulting variational free energy or negative evidence lower bound (ELBO) becomes

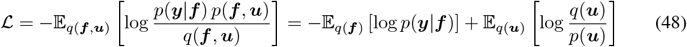

where the first term is the variational expectation of the negative log likelihood with observations ***y***, and the last term is the Kullback-Leibler (KL) divergence of a multivariate normal given by

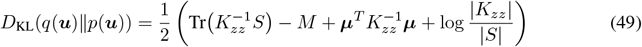

which does not involve the inverse of the full covariance matrix *K*_*xx*_ as for standard GP inference. Minimizing ℒ can then be interpreted as finding the inducing points that optimally summarize the training data ***y***. The variational expectation term involves the sparse posterior

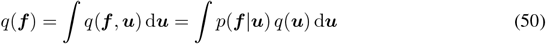

which is another multivariate normal with moments

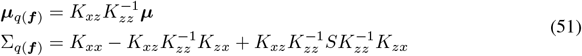

For predictions at new locations {***x***_∗_ **}**, we obtain the posterior predictive distribution by replacing training locations 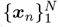 in the sparse posterior above. The decoupling of the data ***y*** from the posterior distributions amortizes the variational inference, which allows mini-batching and hence scalability to large data [16]. Overall, this gives the sparse variational Gaussian process (SVGP).

##### B.1.3 Posterior sampling and Matheron’s rule

Conventionally, one samples from conditionals of multivariate Gaussian distributions by computing the moments and using the Cholesky decomposition. However, this approach comes with 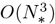 computational complexity, which becomes prohibitive for evaluating posterior function samples at many locations. An alternative to directly working with distributions is the idea of pathwise sampling or Matheron’s rule [37]. For random vectors ***a*** and ***b*** distributed as a multivariate Gaussian we have

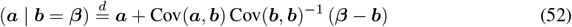

where we have now expressed a procedure involving manipulations of samples rather than distributions. The validity follows from the linearity of the transformation and noting the resulting moments match the posterior Gaussian distribution. In particular, ***a*** and ***b*** are sampled independently.

###### Sparse posteriors

Sparse Gaussian process posteriors are of the form Eq. 50. Matheron’s rule applies to sampling from *p*(***f*** |***u***) with given ***u***. In the sparse posterior, we can interpret the expression as drawing ***β*** ∼ *q*(***u***) and then conditioning *p*(***f*** |***u*** = ***β***). This gives us the posterior distribution written as manipulations on a prior sample *f* (·) ∼ 𝒢 𝒫 (0, *k*(·, ·))

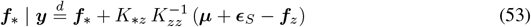

where ***f***_∗_ and ***f***_*z*_ is the same prior sample evaluated at test and inducing point locations, respectively.

###### Pathwise conditioning

From the above, we note that the computational bottleneck is due to sampling the prior ***f***_∗_ at test locations. To work around this, we change our perspective from the function space to the Fourier domain and sample the prior using random Fourier features (see below)

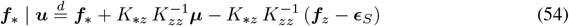

with ***ϵ***_*S*_∼ 𝒩 (**0**, *S*). Because Matheron’s rule decouples the sampling procedure, we can change the method for sampling the prior to obtain this decoupled posterior sampling procedure. This reduces the computational complexity from 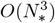 to *O*(*N*_∗_), combining the relative strengths of the functional and Fourier perspectives of GPs [37].

##### B.1.4 Random Fourier features

GPs are conventionally defined in the functional perspective as a prior over functions specified by a covariance kernel function *k*(***x, x***^*′*^). For stationary kernels *k*(***x, x***^*′*^) = *k*(***x*** − ***x***^*′*^), and Bochner’s theorem states that valid stationary covariance functions must have a non-negative Fourier transform 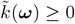. One can interpret this as some probability measure *p*(***ω***) after suitable normalization, and this provides an alternative view on Gaussian process functions as linear combinations of random Fourier features [26]

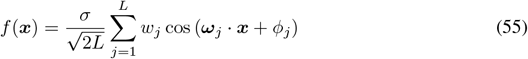

where *σ*^2^ is the kernel variance, *w*_*j*_ ∼ 𝒩 (0, 1), ***ω***_*j*_ ∼ *p*(***ω***) and *ϕ*_*j*_ ∼ 𝒰 (0, 2*π*). For *L* → ∞, this will tend to exact function draws from the GP prior. Note for a product kernel as used in this work, *p*(***ω***) will also be a product of individual kernel factor probability measures.

###### Periodic kernels

For periodic kernels

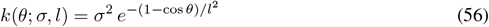

we use the following series expansion [34]

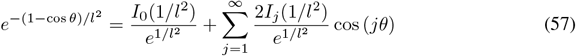

and we note the corresponding probability measure for random Fourier features is discrete. We sample a discrete variable from the truncated series after normalizing the terms

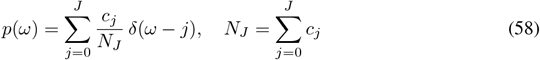

Due to the discrete nature of the probability measure, sample functions are not directly differentiable as we can no longer rely on the reparameterization trick. We can work around this with importance sampling [5], yielding generalized random Fourier features

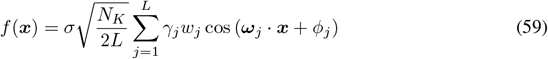

where now ***ω***_*j*_ ∼ *p*_ref_(***ω***) from a reference distribution where the discrete spectrum factors are not sampled from in a differentiable manner, and with importance sampling weights

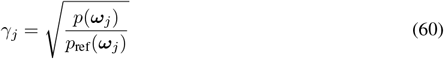

By taking *p*_ref_(***ω***) = *p*(***ω***) we obtain *γ*_*i*_ = 1 but non-zero gradients for each factor on the backward pass. This is achieved by stopping the automatic differentiation for discrete density factors of *p*_ref_(***ω***).

##### B.1.5 Whitened posteriors

The variable transform ***v*** = *L*^−1^***u*** leads to a whitened prior *p*(***v***) = 𝒩 (**0**, *I*). Instead of parameterizing *q*(***u***) = 𝒩 (***μ***, *S*), we can parameterize *q*(***v***) = 𝒩 (***ν***, *W*) linked by the same variable transform. This variational parameterization generally leads to improved conditioning of the inference problem [1], and we implement SVGPs in all models of this paper with the whitened variational parameterization. The ELBO now involves the KL divergence between the variational posterior and the unit normal distribution. Matheron’s rule takes the form with ***ϵ***_*W*_ ∼ 𝒩 (**0**, *W*)

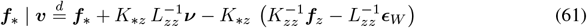

#### B.2 General variational inference framework

##### B.2.1 Constructing probabilistic models

Constructing the baseline models for this paper with GPs involves different probabilistic model structures specific to each model that are described below. However, the inference framework presented for each model is a special case of general probabilistic programming with stochastic variational inference [4, 12, 19] with SVGPs. From this perspective, the negative ELBO generally consists of two parts

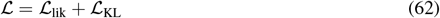

with ℒ _lik_ the variational expectation of the likelihood given the GP posteriors, and ℒ _KL_ the KL divergences of each individual GP component.

##### B.2.2 Data scaling

Our method builds on sparse variational Gaussian processes, and hence inherits the convergence properties and data scaling from such models analyzed in previous work [7, 16, 25]. In this study, our synthetic validation experiment involves 2D inputs ***x***_*t*_ with a million time points, and we are able to accurately recover the ground truth (Fig. 2). Real data has either 1D or 3D *x*_*t*_ and *>* 1 million time points. Based on the synthetic experiment, this suggests that we are in the regime of sufficient data. Note that the total number of GP input dimensions includes the *K* = 4 lagging ISI dimensions for all models.

#### B.3 Rate-rescaled renewal processes

##### B.3.1 Relation to time-rescaling and modulated renewal processes

Note that rate-rescaling for inhomogeneous renewal processes in Eq. 5 is a special case of time-rescaling when applied to the CIF of the rate-rescaled renewal process. As the CIF of renewal processes is related to the ISI distribution via Eq. 22, we can apply change of variables with Eq. 5 as the transformation. Using the shorthand notation *r*(*t*) = *r*(***x***_*t*_), we obtain

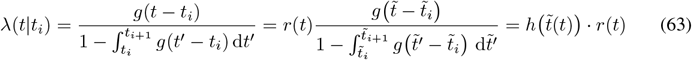

where in the last equality, we wrote in analogous terms to modulated renewal processes Eq. 4. We see that the rate *r* plays a similar role as the modulator *ρ*, but in addition it also affects the hazard function argument via the rate-rescaling integral 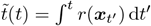. This overall gives

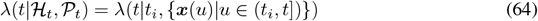

which is a covariate path history dependence rather than an instantaneous dependence *λ*(*t* | ***x***_*t*_, …) like other models in this paper, though in a highly restricted manner through the rate function *r*(***x***). Note we parameterize *g*(∆) such that the mean is always one (Appendix A.3), which allows *r*(*t*) to be interpreted as the instantaneous firing rate of a neuron.

##### B.3.2 Time discretization

In discrete time settings, the transformation applied in practice is discrete rate-rescaling [15]

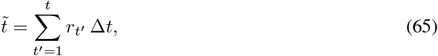

Note that time discretization introduces biases in the KS test rate-rescaled renewal process models [13, 15], as the quantiles are computed using the cumulative density function of continuous-time renewal densities. In particular, very small ISIs in the rescaled time domain will be mapped onto the same time bin in the real time domain if the interval is smaller than the bin size. For our nonparametric non-renewal process model, this bias is not an issue due to the flexibility of the CIF function.

##### B.3.3 Generative model structure

###### Joint versus marginal posterior samples

The structure in the ℒ _lik_ term induced by discrete rate-rescaling Eq. 65 requires us to draw full posterior function samples, as the joint likelihood does not factorize across time steps *p*(***y***| ***f***) ≠*p*(*y*_1_ | *f*_1_) · · · *p*(*y*_*T*_ | *f*_*T*_). The posterior rate function samples are then individually integrated over to perform rate-rescaling. Experiments with joint posterior rate function samples from the GP were significantly slower and less numerically stable failing to fit the data successfully, especially on the hippocampal data. Instead, we use a quasi-MAP approximation where we sample from the marginal posterior rates

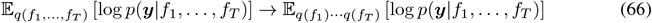

Note that this approximation for ℒ_lik_ no longer leads to a strict negative ELBO. The marginal posterior samples lead to independent noise in the rates, which gives smaller fluctuations in the rescaled times compared to correlated joint posterior samples.

###### Temporal batching and boundary ISIs

As we fit neural population in parallel and temporally batch with fixed batch sizes, each batch will cut off ISIs at the edges. Generally, the likelihood of a given renewal process has boundary terms to account for unobserved spikes outside the temporal range considered, such as incomplete gamma functions in gamma renewal processes [3] or exponential interval terms to approximate the intractable boundary terms [10, 31]. We instead choose to use the rescaled time from the end of the last batch for initializing the rate-rescaling in a given batch, or ignore the first and last boundary terms of the entire spike train in the first and final batches, respectively. As mini-batching cuts the computation graph between batches, we do not back-propagate beyond the batch cutoff and this introduces a small bias to the gradient as. For sufficiently large batch sizes and overall spike train lengths, these boundary effects are negligible.

###### Spike train samples

To sample from the rate-rescaled renewal processes, one first samples from the homogeneous renewal process to obtain spike times 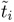. One then applies the transformation shown in Fig. 5A to obtain the real spike times *t*_*i*_, which are modulated according to the rate function *r*(*t*).

#### B.4 Conditional processes

##### B.4.1 Raised cosine spike-history filters

The classical raised cosine basis [35] is defined by

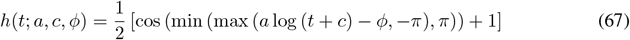

where we can build filters as linear combinations

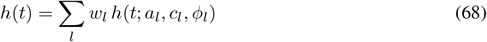

The parameters *a, c* and *ϕ* are fixed while the weights *w* are learned.

##### B.4.2 Nonparametric spike-history filters

Recent work [11] has explored modeling spike-history filters in a nonparametric manner. One can use a Gaussian process with suitable properties for neural data to infer the filter shape in a probabilistic manner. The decaying squared exponential (DSE) kernel encodes suitable inductive biases similar to the ones discussed for our time warping procedure (Eq. 12)

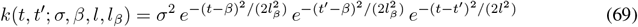

where *β* and lengthscale *l*_*β*_ control the non-stationary aspects of the kernel. Note for *l*_*β*_ →∞, we recover the stationary squared exponential kernel. The GP mean function used is the same as in Eq. 14, now placed on the spike-history time

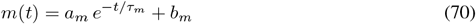

##### B.4.3 Generative model structure

###### Hierarchical GPs

The likelihood term ℒ_lik_ now involves performing a 1D convolution on the spike train. For the nonparametric filters, we sample a filter function *h*(*t*) from the GP posterior and use this same sample throughout the spike train convolution. To handle the resulting intractable hierarchical model, SVGPs are used within the general variational inference framework discussed in Appendix B.2. Note this involves adding an additional KL divergence term of the spike-history filter GP to ℒ_KL_. The parametric raised cosine basis filters simply use point estimates of their parameters, which are treated as hyperparameters of the likelihood.

###### Spike train samples and temporal batching

Temporal batching is straight-forward here as the point process likelihood factorizes across time given the CIF, which we can compute if we condition on the past spike train window. Related to this, we can generate spike trains from this model by using an initial spike-history segment and sampling autoregressively using the CIF.

#### B.5 Bayesian nonparametric non-renewal processes

##### B.5.1 Time warping

For time warping in Eq. 12, we set *τ*_*w*_ as the empirical mean ISI from the training dataset. The time-warped mean function Eq. 14 becomes in the warped domain

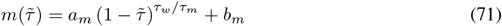

##### B.5.2 Computing conditional ISI distributions and tuning curves

To compute the conditional ISI distribution Eq. 15, we need to compute the integrals appearing in Eq. 21, now conditioned on inputs denoted by *λ*(*t*| …), to obtain a normalized conditional ISI distribution. We define the integral Λ(*t, t*_*i*_| …) which we transform into the warped time domain

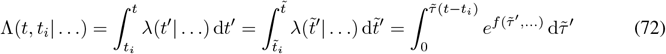

where for 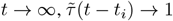. This then allows us to write the normalized Eq. 15

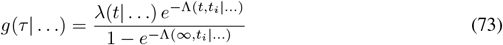

The integrals are evaluated using rescaled Gauss-Legendre quadrature points, as the integral limits cannot exceed [0, 1] using the time warping transform. Standard Gauss-Legendre quadratues is defined for the interval [0, 1], and for smaller intervals we linearly scale the quadrature locations while multiplying the weights by the inverse scaling factor. Note the normalization factor in the denominator is independent of *τ* and simply uses the standard Gauss-Legendre quadrature method.

For a given set of target values *τ*_∗_ where we want to evaluate *g*(*τ*), we need to compute the integrals at a different set of rescaled quadrature points for each target value. This requires us to evaluate a single GP posterior sample 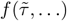 at many locations, which we achieve using pathwise conditioning (see Appendix B.1 for details). In particular, to compute samples of *g*(*τ*) we need to use the same posterior sample for both the numerator and denominator in Eq. 73.

We generally have many *τ*_∗_ values that may only be slightly different, which leads to a large number of rescaled quadrature point locations that are very close to each other. To make this computationally more efficient, we compute the GP posterior samples 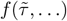 on a regular grid [0, 1] with *G* grid points and store this as a buffer of pre-computed posterior sample values. We then use linear interpolation to obtain 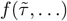 at all the required quadrature locations. Overall, this reduces the required GP posterior function sample evaluations to the *G* grid locations as well as the standard quadrature locations for the normalization factor in Eq. 73. Using a fine enough grid will lead to more accurate values from the interpolation at the cost of a computing and storing a larger buffer.

The moments of *g*(*τ*) can be computed by using an outer loop of rescaled Gauss-Legendre quadratures similar to above with the integrand as given in Eq. 16. Tuning curves of the moments as shown in Fig. 3F and Fig. 4F were computed from joint tuning curve samples, which require evaluating the same posterior ISI distribution sample conditioned on different covariate values. Obtaining all required GP posterior function sample evaluations in a single call would lead to out-of-memory issues, and instead we achieved this using multiple GP posterior function sample evaluation calls while keeping the same pseudo-random number generator key.

##### B.5.3 Generative model structure

Sampling from the generative model. Since the history dependence is now contained in the input covariates **∆**, inference is the same as for inhomogeneous Poisson point processes when conditioned on the full input, and the corresponding loss Eq. 18 is naturally amenable to temporal mini-batching. Note that time warping is a feedforward transformation on the input and hence does not affect temporal batching.

#### B.6 Code

The code is written in Python and utilizes JAX [6] for performing automatic differentiation and numerical optimization for model fitting. All models were implemented from scratch, and we utilize equinox [17] for maintaining readability and elegance of the code. In our code base repository, we provide the code and scripts for reproducing all results in this paper.

### C Further details on experiments

#### C.1 General information

##### Vectorization across neurons

Even though our model is designed for single neuron spike trains, we fit neural populations in parallel by vectorizing over all neurons. Each neuron has its own copy of model (hyper)parameters. This implementation also allows straight-forward incorporation of shared latent variables for modeling neural correlations.

##### Optimization hyperparameters

We use the Adam [18] optimizer, where the learning rate is set to 10^−2^ at the start and annealed down to 10^−4^ with a decay factor per epoch of 0.998 for synthetic experiments and 0.98 for real data experiments. The stopping criterion for model fitting is when 100 epochs have elapsed and the average training loss in each epoch has decreased less than 10^0^ for synthetic and rat data experiments, and 10^1^ for mouse data experiments. We also set a maximum number of 3000 epochs, though in practice most experiments finish before this maximum is reached. We use a temporal batch size of 10000 time points (10 s) for all models except rate-rescaled models where we use a batch size of 30000 time points (30 s). Experiments are run with single floating point precision, and a jitter of 10^−6^ is used to stabilize the Cholesky decomposition in GPs.

##### Parameter constraints

Kernel lengthscales, variances and other positive parameters are parameterized as unconstrained parameters pushed through softplus or exponential transformations. Variational distribution covariances are parameterized with a lower triangular matrix where we enforce positive diagonal elements at least as big as the jitter value used. Shape parameters for renewal densities were additionally constrained to be *α* ≥10^−1^ for gamma and *μ* ≥10^−5^ for inverse Gaussian renewal densities to ensure numerical stability.

##### Variational expectation evaluation

During training, we use 20 Gauss-Hermite quadrature points to evaluate the variational expectation for all models except rate-rescaled renewal processes. There, we use one Monte Carlo sample from the marginal posterior of the GP rate function to estimate the variational expectation of the likelihood (see Appendix B.3 for a discussion on the GP marginal versus joint posterior sampling for rate-rescaling). For evaluating models on test data, we use 50 Gauss-Hermite quadrature points to evaluate the variational expectation terms and 10 Monte Carlo samples for the rate-rescaled renewal processes.

##### Inducing point initialization

Randomly initialized inducing points by picking random locations without replacement from a *D*-dimensional regular grid on the input covariate space. Inducing point locations are learned as part of the hyperparameters. Note that sparse GPs can suffer from numerical instabilities when some inducing points get very close to each other and cause problematic conditioning numbers, which tend to arise when the total number of inducing points is high [32].

##### Conditional process hyperparameters

For the raised cosine filters in conditional processes (Appendix B.4), we follow [35] and use a 150 ms window with the parameters *a* = 4.5, *c* = 9, and 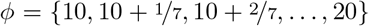 for defining our raised cosine basis consisting of 8 basis functions (Fig. 5B top). Note these parameters are fixed and not optimized during training. The nonparametric conditional Poisson process models spike-history filters using the GP defined in Appendix B.4. Hyperparameters of the kernel are learned during model fitting. We use 6 inducing points initialized uniformly along the 150 ms time lag axis of the spike-history filter.

##### Computing resources and reproducibility

Experiments were ran for training each model or performing analysis of fitted models on single NVIDIA GeForce RTX 2080 Ti GPUs with 11 GB of memory. Each model fitting run can take up to 6 hours on this hardware, but is generally a lot shorter due to the termination criterion. Analysis scripts of fitted models were also run on the same hardware and can take up to 8 hours for the most compute-intensive tasks. The code provided contains a bash script with the exact commands and random seeds that were run to generate the results in this paper. Datasets were taken from the public online database https://crcns.org/, with specific instructions on which datasets to use and preprocessing scripts provided in the code repository.

##### Neural data preprocessing

Electrophysiological neural recordings typically provide spike times at higher temporal resolution than animal behavior, the latter being inferred from video recordings of the animal. To match the input covariates and spike trains in our discrete time point process, we upsample the behavior using Akima interpolation, which suffers less from oscillations when sharp changes are present in the time series [2]. This leads to data sampled at 1 ms regular intervals. From the spike trains, we also compute lagging ISIs at this temporal resolution. Note they include a section at the start of the neural recording where we have undefined lagging ISIs of higher order. To obtain the dataset segment we use, we cut out the section at the start until we have defined lagging ISIs up and until order *K* = 4. We also correct for duplicate spike counts in the 1 ms bins, which are artifacts of spike sorting errors. The number of such duplicate spike counts was minimal, as we also selected for cells which had a fraction of 2 ms refractory period violations less than 5%.

#### C.2 Validation on synthetic data

For the synthetic population shown in Fig. 2A, the densities used were (each column, top to bottom) the gamma with *α* = {0.5, 1.0, 1, 5}, the log normal with *μ* = {0.5, 1.0, 1, 5}, and the inverse Gaussian with *σ* = {0.5, 1.0, 1, 5} (equations given in Appendix A.3). Though the Poisson case (left middle dark green) stricly has no past spike dependence in the CIF (i.e. independent of *τ*), the time warping shape requires nonlinear tuning of the CIF to 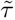.

We use 16 GP inducing points for baseline models and 48 for our NPNR process. Firing rates and CVs from the NPNR model were computed using 30 posterior samples. For computing the condition ISI distributions, we used *G* = 1000 grid points and 100 Gauss-Legendre quadrature points (see Appendix B.5).

#### C.3 Neural data

Violin plots of KS *p*-values in Fig. 3A and Fig. 4A show kernel density estimates using the Silverman method [29]. Instantaneous firing rates and CVs as well as tuning curves of the NPNR model are computed using 50 posterior samples. For computing the condition ISI distributions, we use *G* = 1000 grid points and 100 Gauss-Legendre quadrature points (see Appendix B.5). For calculating the coefficient of determination shown in Fig. 3E and Fig. 4E, we use linear regression on the instantaneous rate and CV estimates (posterior means)… Standard SVGP regression [16] using 8 inducing points with a squared exponential kernel is applied to the log rate and CV, where the logarithmic transform helps to account for the non-stationary lengthscale of CV-rate scatter cloud shapes along the rate axis.

##### Neuron selection criteria

For selecting putative neurons in the neural recordings, we use the mutual information (MI) per spike, tuning curve coherence and tuning curve sparsity measures [38], which are computed from simple histograms of spiking activity and occupancy over binned relevant covariates. For each bin *i*, we compute the number of time steps *T*_*i*_ when the covariates ***x***_*t*_ are located in that bin. We also compute the total number of spikes *N*_*i*_. Now we can compute the average rate *r*_*i*_ and relative occupancy distribution *P*_*i*_

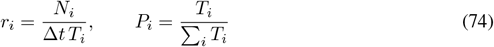

with time bin size ∆*t*. From the rate map *r*_*i*_, we obtain a smoothed rate map 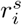 which is given by the convolution of a smoothing kernel with *r*_*i*_. Using these quantities, we get the tuning measures

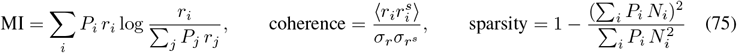

where ⟨… ⟩ denotes the average and *σ*_…_ is the standard deviation. Note the coherence is identical to the Pearson correlation coefficient between the smoothed and original histogram rate maps.

##### Mouse head direction cell data

We pick the wake portion in session “140313” of mouse “28” from the mouse dataset [21, 22], which involves a freely moving mouse foraging for food with neural recordings from anterodorsal nucleus post-subiculum. Putative head direction cells are selected based on refractory violation fraction *<* 2%, head direction mutual information *>* 0.5 bits per spike and head direction tuning curve sparsity *>* 0.2. These tuning measures are computed using a binning of head direction into 60 equal bins. Overall, we end up with 1085970 time points for training data. As we have one dimension (head direction) for our behavioural covariates, we use 8 GP inducing points for baseline model GPs and 40 for our NPNR process (additional *K* = 3 lagging ISI dimensions plus the *τ* dimension).

##### Rat place cell data

We pick session “ec014.468” from the rat dataset, which involves the animal running along a linear track of 250 cm with neural recordings from hippocampal area CA1 using silicon-based electrodes [20]. Putative place cells are selected based on refractory violation fraction *<* 2%, joint *x*-*θ* mutual information *>* 0.5 bits per spike and joint *x*-*θ* tuning curve sparsity *>* 0.6, coherence *>* 0.4, minimum number of spikes *>* 900 and more than 5 spikes in the first 100 s of the dataset segment used. These tuning measures are computed using a binning of *x* and *θ* into 40 and 30 equal bins, respectively. The smoothing kernel for computing the coherence is a 2D factorized Gaussian with standard deviations of 2 bins in each dimension. Overall, we end up with 1934126 time points for training data. We use 32 inducing points for baseline model GPs and 64 for our NPNR process (additional *K* = 3 lagging ISI dimensions plus the *τ* dimension). Note that this uses slightly more than 8 inducing points per dimension of ***x***, which we choose to help the model capture potentially more complicated tuning curves relevant to *θ*-phase precession patterns. The GP jitter is increased to 10^−5^ for the nonparametric conditional Poisson process as it is less stable numerically on this dataset.

### D Additional results

#### D.1 Validation on synthetic data

##### Test expected log likelihoods

On synthetic data, the expected log likelihood (ELL) values on test data consisting of 5 different held-out synthetic datasets generated from the same ground truth model are shown in Table 1. This shows that our model outperforms the (conditional) Poisson baselines, while performing comparable to the Gamma renewal process (which is within model class for three of the synthetic neurons) within dataset variability. Overall, these numbers are consistent with the goodness-of-fit results shown in Fig. 2C.

**Table 1:**
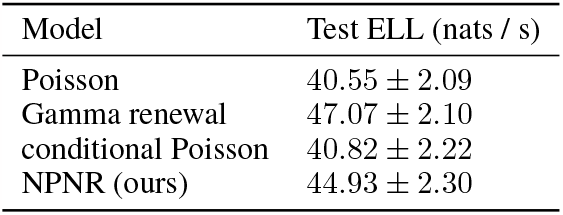
Test expected log likelihoods of models on synthetic data.

| Model | Test ELL (nats / s) |
| --- | --- |
| Poisson | $40.55 \pm 2.09$ |
| Gamma renewal | $47.07 \pm 2.10$ |
| conditional Poisson | $40.82 \pm 2.22$ |
| NPNR (ours) | $44.93 \pm 2.30$ |

##### Automatic relevance determination

The time warping component of our model has a parameter *τ*_*w*_, which sets the time range of *τ* that is mapped into the range of 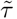 where the stationary GP can meaningfully model temporal fluctuations. In the main text, we propose to look at kernel timescales to determine relevance of lagging ISI dimensions. However, *τ*_*w*_ also interferes with the effect of the kernel timescales on the effective timescale in the *τ* and **∆** dimensions of the log CIF. Indeed, we see that learning *τ*_*w*_ generally leads to loss of interpretability for the GP kernel timescales in Fig. 6A. By fixing it to the empirical mean ISI for each neuron, we obtain a suitably normalized time warping scheme where the GP kernel lengthscales are identifiable with temporal fluctuations of the log CIF, and hence the relevance of lagging ISI dimensions.

##### Rate map estimation

Visually, the inferred rat maps for the Poisson, rate-rescaled Gamma and nonparametric non-renewal (NPNR) models are quite close to the ground truth. The conditional Poisson model has estimated rate maps that do not match the truth, showing the complicated effects of the spike-history filter on the effective firing rate and smoothness of the rate profile.

#### D.2 Neural data

##### Head direction cell data

We plot tuning curves in Fig. 8 for the rate and CV for all neurons in our processed dataset. Each time point of posterior mean estimated instantaneous rates and CVs is plotted as a dot in Fig. 7, where we only show the estimates statistics per 10 ms interval.

##### Place cell data

We perform the same analysis as for head direction cells, but now the tuning curves are 2D maps over *x*-position and *θ*-phase. In addition, the animal can run left-to-right or vice versa, which we model by including the head direction covariate of the animal. Tuning curves in Fig. 10 and Fig. 11 are then computed conditioned on the head direction for each run direction. Each time point of posterior mean estimated instantaneous rates and CVs is plotted as a dot in Fig. 9, where we only show the estimates statistics per 10 ms interval.

**Figure 10:**
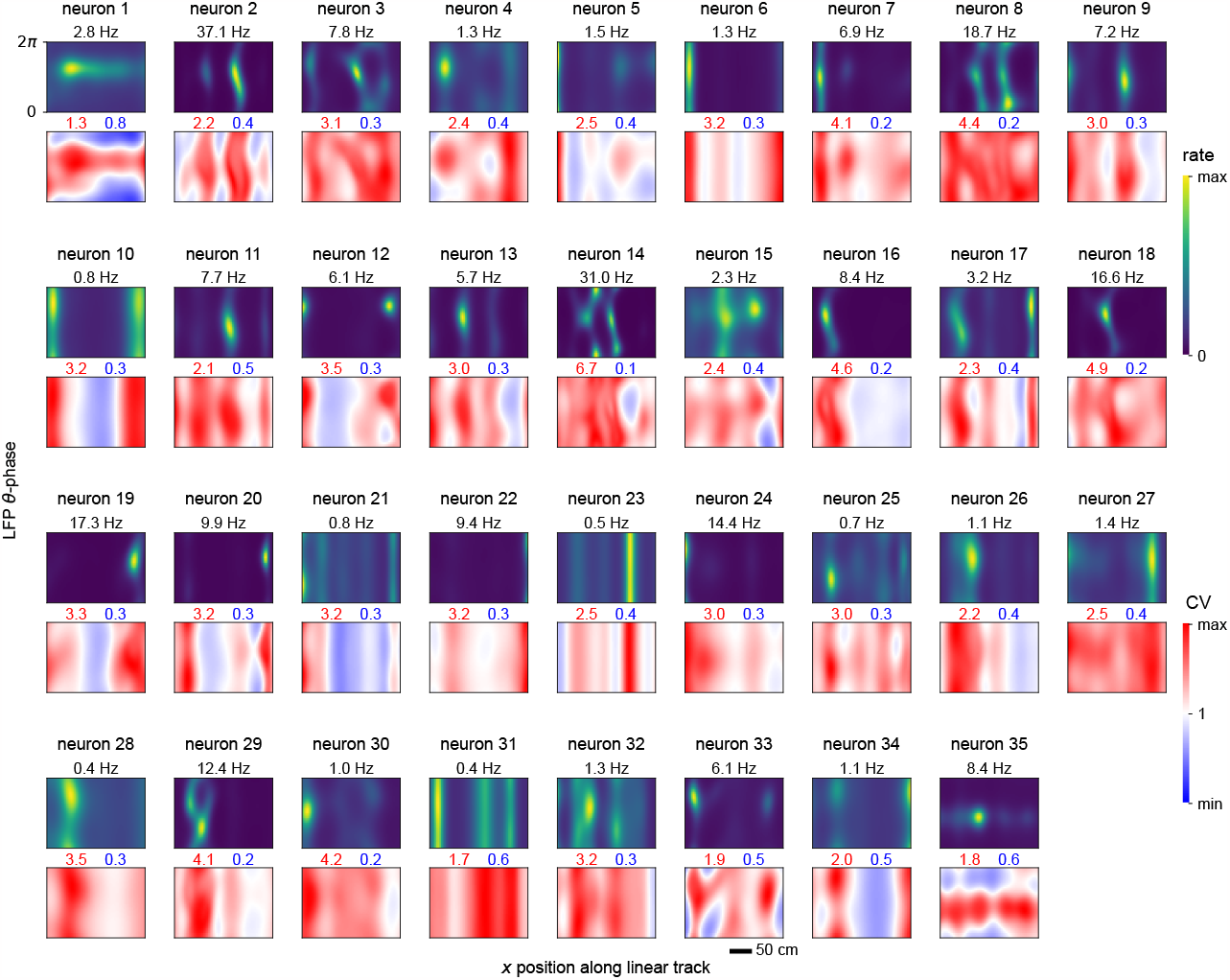
Tuning curves of place cells for rat running left-to-right. Heat maps show posterior mean values of spike train statistics computed from the conditional ISI distribution samples.

**Figure 11:**
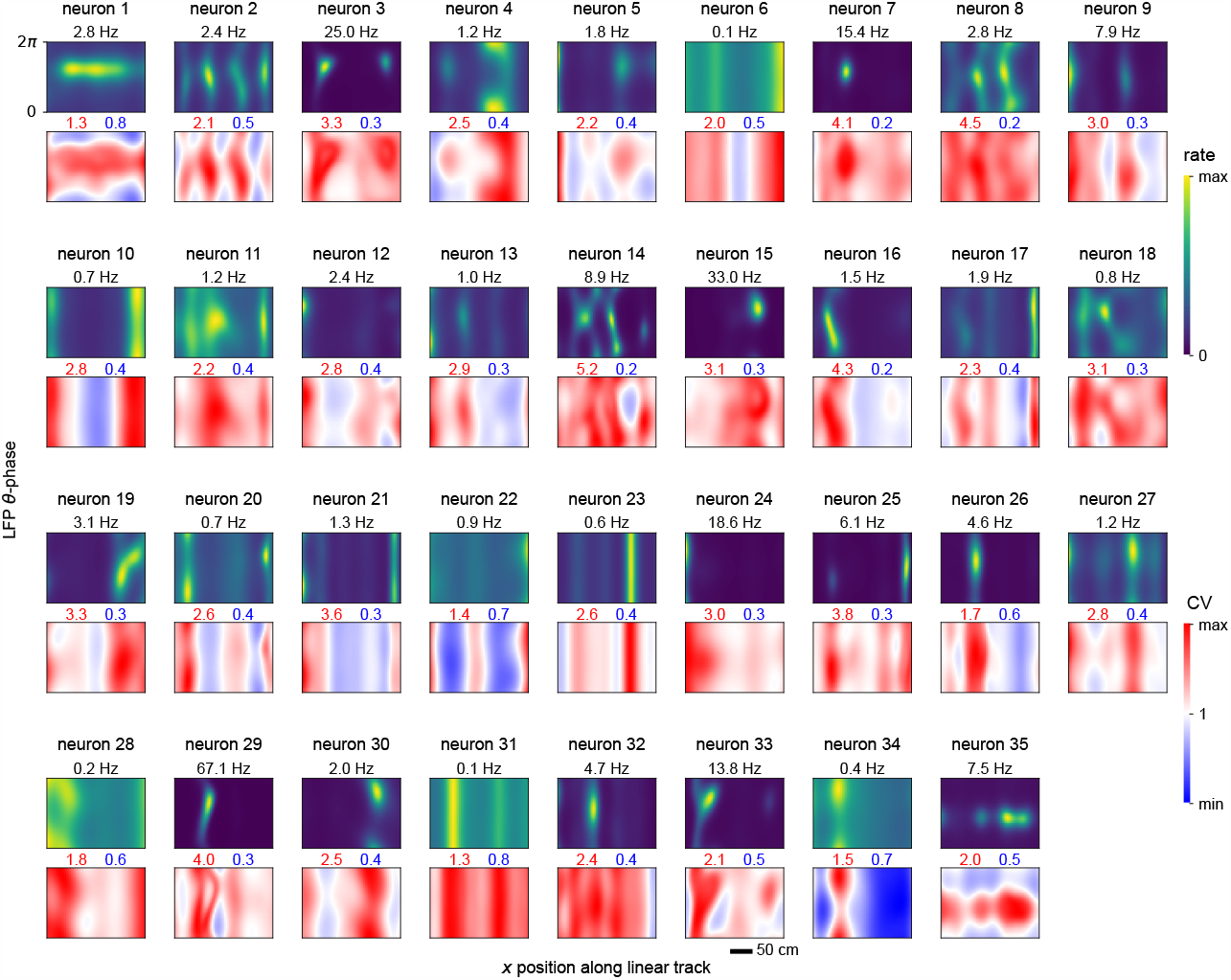
Tuning curves of place cells for rat running right-to-left. Heat maps show posterior mean values of spike train statistics computed from the conditional ISI distribution samples.

**Figure 12:**
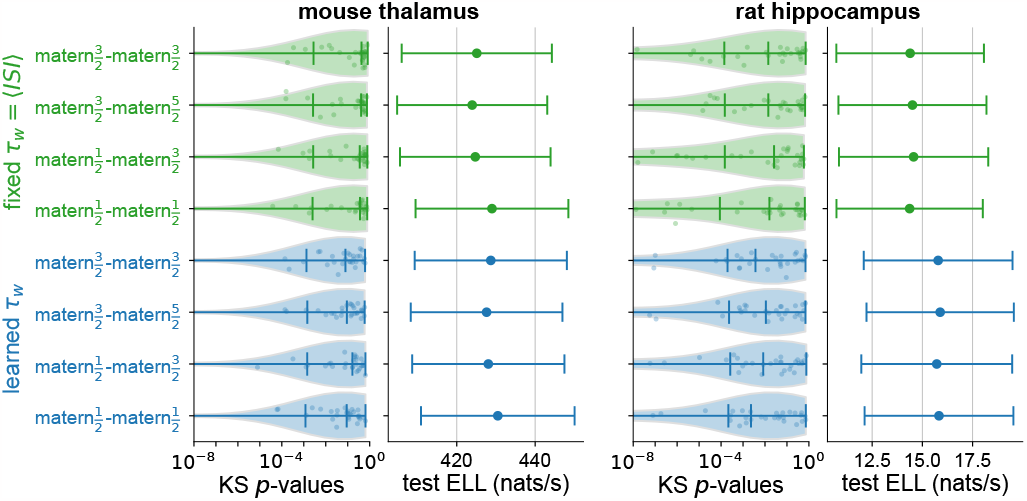
Comparison of kernel selection for the non-renewal process. We show measures of model fits to real data similar to the model in the main text for various temporal kernel choices in the format 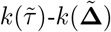, and fixed or learned time warping scale *τ*_*w*_ (green or blue, respectively).

##### Temporal kernel selection

We compare Matérn kernels of different order to use as temporal kernels in our model, as well as fixing the time warping parameter *τ*_*w*_. From Fig. 12, we see that the differences in performance are small. Learning *τ*_*w*_ generally bumps up the predictive performance, but has mixed effects on the KS *p*-value distributions. The choice made in the main paper (Matérn-^3^*/*_2_ for both 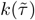 and 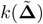) gives good performance in both predictive power and KS *p*-values across both neural datasets.

#### D.3 Additional ISI statistics and variability measures

Our method captures joint lagging ISI statistics from data into *g*(*τ* | **∆**, …). This provides a more general approach to computing correlations between consecutive rate-rescaled ISIs [23]. As a specific example, the local coefficient of variation [28] can be computed form our model as

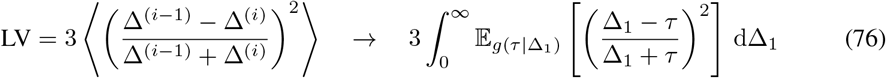

which we can perform using Gauss-Legendre quadratures and time warping similar to Eq. 16, but using a 2D quadrature grid as we now deal with a double integral.

## Footnotes

1 Code available at https://github.com/davindicode/nonparametric-nonrenewal-process

2 Note the simple summation over *n* in Eq. 18, as our model does not capture neural correlations without introducing latent covariates [48]. In other words, in its current form, our model treats neurons as independent.

